# Induction of mitochondrial heat shock proteins and mitochondrial biogenesis in endothelial cells upon acute methylglyoxal stress: Evidence for hormetic autofeedback

**DOI:** 10.1101/2021.11.30.470545

**Authors:** Ruben Bulkescher, Thomas Fleming, Claus Rodemer, Rebekka Medert, Marc Freichel, Matthias Mayer, Julia Szendroedi, Stephan Herzig, Johanna Zemva

**Author notes:** Corresponding author. E-Mail (JZ).

## Abstract

Increased metabolic flux produces potentially harmful side-products, such as reactive dicarbonyl and oxygen species. The reactive dicarbonly methylglyoxal (MG) can impair oxidative capacity, which is downregulated in type 2 diabetes. Heat shock proteins (HSPs) of subfamily A (Hsp70s) promote ATP-dependent processing of damaged proteins during MG exposure which also involve mitochondrial proteins. Since the protection of mitochondrial proteins could promote higher production of reactive metabolites due to increased substrate flux, tight regulation of HspA-mediated protein handling is important. We hypothesized that stress-inducible HspAs (HspA1A/HspA1B) are pivotal for maintaining mitochondrial biogenesis during acute MG-stress. To analyze the role of stress-inducible HspA1A/HspA1B for maintenance of mitochondrial homeostasis during acute MG exposure, we knocked out HSPA1A/HSPA1B in mouse endothelial cells. HSPA1A/HSPA1B KO cells showed upregulation of the mitochondrial chaperones HspA9 (mitochondrial Hsp70/mortalin) and HspD1 (Hsp60) as well as induction of mitochondrial biogenesis upon MG exposure. Increased mitochondrial biogenesis was reflected by elevated mitochondrial branching, total count and area as well as by upregulation of mitochondrial proteins and corresponding transcription factors. Our findings suggest that mitochondrial HspA9 and HspD1 promote mitochondrial biogenesis during acute MG stress, which is counterregulated by HspA1A/HspA1B to prevent mitochondrial overstimulation and to maintain balanced oxidative capacity under metabolic stress conditions. These data support an important role of HSPs in MG-induced hormesis.

## 1. Introduction

During increased metabolic flux, reactive side-products are inevitably produced, such as reactive dicarbonyls from glycolysis or reactive oxygen species (ROS) from oxidative phosphorylation. The concept of hormesis, meaning that low doses of a substance are beneficial whereas high doses are toxic, is well defined for a broad spectrum of metabolites (1). Next to physical and chemical agents, intrinsic metabolites were shown to induce hormetic reactions in model organisms but also in humans. Accordingly, beneficial effects achieved by physical training were diminished by simultaneous supplementation of antioxidants, possibly via scavenging of ROS thereby inhibiting ROS-induced mitohormesis (2). Moreover, mitochondrial adaptation to increased substrate flux and ROS-production was shown to counteract the development of steatosis and steatohepatitis in non-alcoholic fatty liver disease (NAFLD) (3). The reactive dicarbonyl methylglyoxal (MG) was described to induce a hormetic reaction by increasing cell survival to different stressors in yeast cells (4) and healthy aging in *C. elegans* (5). However, it stays unclear, if MG-induced hormesis is also relevant in the mammalian system and if it also involves mitochondrial capacity.

Under physiological conditions, MG is predominantly formed as a spontaneous by-product from the intermediates glyceraldehyde 3-phosphate and dihydroxyacetone phosphate (triosephosphates) during glycolysis (6, 7). MG can react with and thereby modify proteins, lipids, and DNA, resulting in the formation of advanced glycation endproducts (AGEs) (8). Regarding protein modifications, the most abundant adduct is the modification of arginine residues, also called hydroimidazolone or MG-H1 (7, 8). Posttranslational modifications of proteins lead to conformational changes, impaired function, and increased risk of aggregation. MG and AGEs have been shown to be involved in the development and progression of diabetic complications, such as diabetic nephropathy (9-12) or cardiovascular disease (13, 14). AGEs also contribute to reduced oxidative capacity, which is a hallmark of diabetic complications, and has been shown to relate to insulin resistance and NAFLD (15). Also, MG has been linked to mitochondrial dysfunction, which is supported by the identification of MG-modifications within the mitochondria (16-19). Micro- and macrovascular damage is one of the major drivers of diabetic complications and is characterized by endothelial disruption. However, the exact mechanisms linking increased AGE formation and mitochondrial dysfunction in endothelial cells is still to be unraveled.

Heat shock proteins (HSPs) are critical for diverse cellular housekeeping functions, including refolding misfolded or damaged proteins (20, 21). Modified proteins cannot be solely removed by molecular chaperones or HSPs but require further components of the protein quality control (PQC), like the proteasome or autophagy. However, HSPs can prevent the aggregation of modified proteins and play a central role in protecting cells from proteotoxic stress (22). The HspA/Hsp70 family comprises ATP-dependent chaperones with several members, sharing similar structure and function (23). The stress-inducible members HspA1A and HspA1B are predominantly expressed in the cytosol, highly homologous and can fully compensate each other reflecting mutual regulation of HSPs. The constitutively expressed cytosolic member HspA8 or Hsc70 is essential for housekeeping functions (24). HspA9, also called mtHsp70 or mortalin, is a compartment-specific member and exclusively expressed in the mitochondria. While mitochondrial HSPs can be upregulated upon the mitochondrial unfolded protein response (mtUPR) (25) or during mitochondrial biogenesis (26), complete knockout of HspA8 or HspA9 is lethal (27). In the absence of ATP, HspA family members strongly bind to misfolded proteins and are released upon ATP binding to the N-terminal region. ATP-binding, hydrolysis (for example by members of the Dnaj/Hsp40 family) and removing of ADP is facilitated by co-chaperones or small heat shock proteins (28). This cycle of binding and release of the misfolded protein is repeated until complete refolding is achieved (29). Therefore, regulation of the cellular energy level and of mitochondrial biogenesis are critical for efficient functioning of the PQC. Furthermore, mitochondrial dynamics including fission and fusion are essential for mitochondrial PQC and homeostasis (30). Fusion contributes to mitochondrial elongation and maintains cellular oxidative capacity (31) whereas fission can induce a highly fragmented mitochondrial network resulting in decreased ATP production (32). Therefore, the cellular ATP-levels are highly dependent on the overall mitochondrial mass and interconnection. However, it is still unclear how the cell regulates or compensates increased ATP-demand upon rising proteotoxic stress.

In line with this, patients with type 2 diabetes show decreased levels of the key-regulator of mitochondrial biogenesis Ppargc1a (Peroxisome proliferator-activated receptor gamma coactivator 1-alpha) (33) and Ppargc1a responsive genes involved in oxidative phosphorylation (34).

The HspA family is found to be involved in many disease conditions, including type 1 and type 2 diabetes (T2D) (35, 36). Single nucleotide polymorphisms (SNPs) of HSPA1A/HSPA1B have been linked to the development of diabetic nephropathy in T2D (37-39). Furthermore, decreased expression of skeletal HspA1A/HspA1B has been shown to be associated with insulin resistance (40, 41). A negative correlation with age has been reported for HspA1A/HspA1B in T2D (42). Apart from HspA1A/HspA1B, the mitochondrial expressed HspD1 (Hsp60) and the small HspB1 (Hsp27) have also been linked to micro- and macrovascular complications in T2D (43-45). So far, the role of HSPs in preventing accumulation of MG-modified proteins and preserving mitochondrial homeostasis under acute MG stress is still unknown.

To understand the role of HSPs for maintenance of mitochondrial homeostasis during rising proteotoxic stress, and the mutual regulation of different HSPs, we analyzed the effect of acute MG-exposure on mitochondrial biogenesis and HSPs in mouse cardiac endothelial cells (MCECs). We hypothesized that this would aggravate accumulation of MG-modified proteins and disrupt mitochondrial homeostasis upon acute MG-stress. Furthermore, we questioned if induction of MG-driven hormesis would be diminished in the absence of stress-inducible HSPA1A/HSPA1B, as the yeast homologue of HSPA1A/HSPA1B was shown to be a key mediator of the MG-induced defense response (4).

## 2. Material and Methods

### 2.1 Cell culture

An immortalized mouse cardiac endothelial cell (MCEC) line was obtained from Cellutions Biosystems (#CLU510). Cells were cultivated in DMEM (Gibco, #31885023) supplemented with 5% FCS (Sigma, #F4135), 1% penicillin (10,000 Units/ml) (Gibco), 1% streptomycin (10 mg/ml) (Gibco), 1% amphotericin B (250 μg/ml) (Gibco) and 1 mM HEPES (Gibco) in a humidified atmosphere at 37 °C and 5% CO_2_. Cells were grown to full confluency and then passaged with 0.05% Trypsin (Gibco) in gelatin-coated (0.5% in PBS for 15 minutes) cell culture flasks. All cell lines were regularly tested for mycoplasma contamination.

### 2.2 Generation of HSPA1A/HSPA1B knockout cell line

1×10^6^ cells were transfected (Neon Transfection System, Invitrogen) with two vectors from Sigma-Aldrich, targeting the two stress-inducible Hsp70 variants Hspa1a (Gene ID: 193740; targeting sequence of the gRNA: TGTGCTCAGACCTGTTCCG) and Hspa1b (Gene ID: 15511; targeting sequence of the gRNA: CGGTTCGAAGAGCTGTGCT). Both vectors contained one of the respective gRNA target sequences, the Cas9 endonuclease gene and a fluorescent reporter gene (GFP for Hspa1a and RFP for Hspa1b). Fluorescence activated cell sorting (FACS) was performed to detect and isolate GFP and RFP expressing cells. Clones were cultured and genome, mRNA and protein analysis were performed to confirm successful knockout of HSPA1a/HSPA1B. Cell clones AD4 and BE12 were confirmed as full double knockout clones of both genes on the genome, mRNA and protein level. The third clone BH9 was confirmed as a full double knockout clone on the mRNA and protein level.

### 2.3 Methylglyoxal treatment

75,000 cells/cm^2^ were seeded in a gelatin-coated (0.5% in PBS for 15 minutes) cell culture dish (T75-flask or 60mm petri dish or 96-well plate). The next day, cells were washed with PBS and replaced with medium containing 0.1% FCS (assay medium) for 1 hour. Then, methylglyoxal (Sigma-Aldrich) was added to the assay medium to a concentration of 500 µM and added to the cells. Only assay medium was added to the control. For RNA measurements (RT-qPCR and mRNA-Seq) the cells were harvested after 12 hrs, for protein measurements (Fluorescence microscopy and Western blotting) cells were fixed or harvested after 24 hrs.

### 2.4 Immunocytochemistry and Fluorescence microscopy

For immunofluorescent staining, cells were fixed with 4% paraformaldehyde solution for 20 min at RT and permeabilized with 0.1% Triton X-100 for 8 min. Then, blocking was performed with 3% BSA in PBS for 60 min. All primary antibodies were added in blocking buffer in dilutions of 1:50 – 1:300 and incubated overnight at 4 °C. All secondary antibodies were added in blocking buffer in a dilution of 1:1000 and incubated for 2 hrs at RT. Nuclear counterstaining was performed with Hoechst 33342 (#H3570, Molecular Probes) with a working solution of 1 μg/mL in PBS for 1 min 30 sec at RT. Between all steps, the cells were washed three times for 5 min with 1× PBS at RT. A list of all antibodies can be found in the supplementary material (Supplementary material Table S1).

For the acquisition of the images, the automated screening widefield microscope IX81 from Olympus was used, with the ScanR acquisition software. The images were taken with a 60X objective and the following filters: DAPI filter (absorption maximum: 358 nm; emission maximum: 461 nm; for detection of cell nuclei stained with Hoechst 33342); GFP filter (absorption maximum: 395 nm; emission maximum: 475 nm), Cy3 filter (absorption maximum: 550 nm; emission maximum: 570 nm), and Cy5 filter (absorption maximum: 650 nm; emission maximum: 670 nm). Image analysis was performed using the Java-based image processing program ImageJ (http://imagej.nih.gov/ij/) and KNIME software (www.knime.org).

### 2.5 RNA isolation and Reverse-transcription quantitative PCR (RT-qPCR)

RNA isolation was performed with the RNeasy Mini Kit (Qiagen) according to the manufacturer’s protocol. cDNA transcription was performed with the High-Capacity cDNA Reverse Transcription Kit (Applied Biosystems) according to the manufacturer’s protocol. RT-qPCR was performed using PowerUp™ SYBR™ Green Master Mix (Applied Biosystems) on a StepOnePlus™ Real-Time PCR System (Applied Biosystems).

Signals of amplified products were verified using melting curve analysis and mRNA levels were normalized to Hypoxanthine-guanine phosphoribosyl transferase (Hprt, Gene ID: 15452). The fold-changes in gene expression levels were calculated using the ΔΔCt method. Primer sequences used for analyzing mRNA can be found in the supplementary material (Supplementary material Table S1).

### 2.6 MG-H1 clearance assay

24 hrs after treatment (see 2.3), MG-containing assay medium was removed, the cells were washed with PBS three times and assay medium was added to the cells. The plates were fixed at 0, 24 and 48 hrs after MG removal and stained according to 2.4. The plates were measured on an Odyssey DLx Imaging system (LI-COR) and analyzed using Image Studio™ Lite (https://licor.com/bio/image-studio-lite/).

### 2.7 Mitochondrial network analysis (imageJ plugin)

Mitochondrial network analysis was performed with a self-made imageJ plugin/macro. For this, images were acquired as z-stacks of a total of 30 layers with a step size of 200 nm, followed by deconvolution using the Huygens professional software (https://svi.nl/HomePage). Settings that can be modified were image processing (background subtraction), thresholding, size and shape discrimination, binary image processing and mitochondria network branching analysis. The images were first converted to a binary mask and then skeletonized. The skeletonized images were then analyzed regarding number, size, and signal intensity of the particles. For the mitochondrial signal, locations with three or more neighboring pixels were counted as branches/junctions. A more detailed depiction of the complete workflow can be found in the supplementary material (Supplementary Text S2).

### 2.8 Western blotting

20 µg protein of the cell lysates (see 2.8) were mixed with 4x Laemmli buffer and heated to 95°C for 5 min. The separation was done in a Mini-PROTEAN® TGX (Bio-Rad) precasted gel (4–20% acrylamide) at 150V for 75 min. Proteins were transferred on a nitrocellulose membrane in a Trans-Blot® Turbo™ Transfer System (Bio-Rad) and blocked with 5% non-fat dry milk in PBS or protein-fee blocking buffer (Pierce) containing 5% goat serum for 1 hour at RT. When necessary, endogenous biotin sites were blocked with the avidin/biotin blocking kit from Linaris following the manufacturer’s protocol. Membranes were incubated with the primary antibody in blocking buffer at 4 °C overnight, and incubated with the secondary antibody in blocking buffer for 1 hour at RT. Between all steps, the membranes were washed three times for 10 min with 1× TBS-Tween20 (0.1% (v/v)) at RT. The bands were detected on a ChemiDoc imaging system (Bio-Rad) with ECL detection reagent (GE Healthcare) or on an Odyssey DLx Imaging system (LI-COR) and analyzed using imageJ (http://imagej.nih.gov/ij/) or Image Studio™ Lite (https://licor.com/bio/image-studio-lite/).

### 2.9 Statistical analysis

Experimental results are expressed as mean ± standard deviation or, where stated, as mean ± standard error of the mean. Depending on the experimental setup, statistical significance was analyzed using ordinary one-way or two-way ANOVA. The analysis was performed with GraphPad Prism software (https://www.graphpad.com/) and p-values < 0.05 were considered statistically significant.

## 3. Results

### 3.1 MG-H1 accumulation is higher in HSPA1A/HSPA1B KO compared to WT cells upon acute MG-stress

To understand the role of stress-inducible HspA1A/HspA1B for processing of MG-modified proteins and MG-H1 accumulation, we generated HSPA1A/HSPA1B knockout MCECs using Crispr-Cas9 technology as shown in Fig S1. After 24 hrs of incubation with 500 µM MG, HSPA1A/HSPA1B KO cells had higher MG-H1 levels as compared to WT cells (Fig 1A, Fig 1B). To analyze the capacity of both cell lines to clear MG-H1, medium was changed after 24 hrs and MG-H1 concentrations were measured after 24, 48 and 72 hrs. Compared to WT cells, HSPA1A/HSPA1B KO cells initially accumulated higher MG-H1 levels (Fig 1C). However, after changing the medium at 24 hrs to MG-free DMEM, both WT and HSPA1A/HSPA1B KO cells were able to clear MG-H1 adducts to a similar degree, reaching levels close to untreated controls after 48 hrs (Fig 1C).

**Fig 1.**
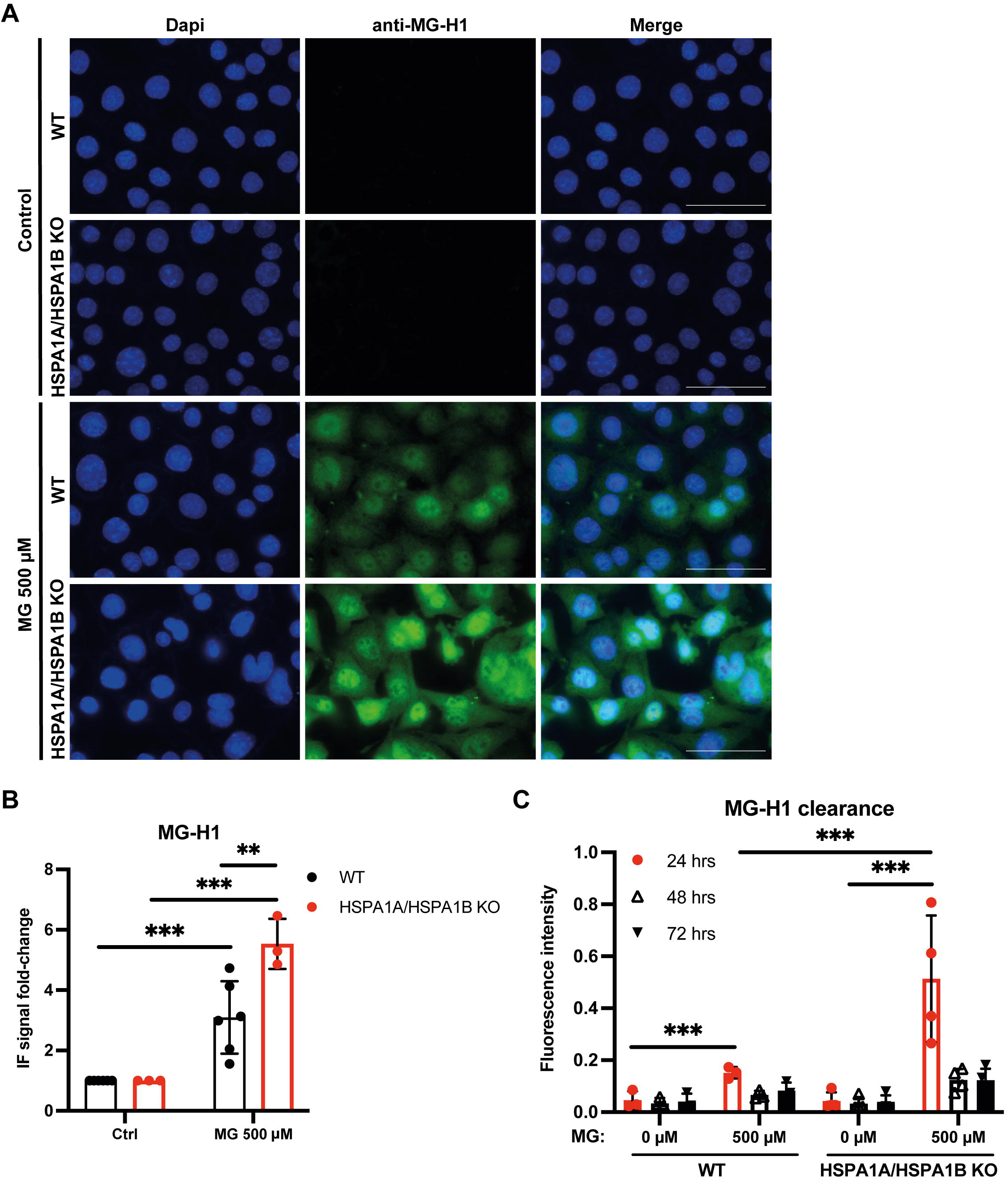
MG-H1 accumulation is higher in HSPA1A/HSPA1B KO compared to WT cells upon acute MG-stress. Immunofluorescence staining with an anti-MG-H1 antibody of WT and HSPA1A/HSPA1B KO cells was performed after MG-treatment with 500 µM for 24 hrs (A) and quantified in (B). (C) MG-H1 clearance was monitored after 24 hrs MG-treatment in HSPA1A/HSPA1B KO vs. WT cells. Results are shown as means ±SD of at least 3 independent experiments. Two-way ANOVA with Šídák’s multiple comparisons test was performed for statistical analysis, *p<0.05, **p<0.01, ***p<0.001.

### 3.2 MG-stress induces mitochondrial heat shock proteins which is counteracted by HspA1A/HspA1B

As the PQC and HSP system is tightly regulated, the effective MG-H1 clearance in HSPA1A/HSPA1B KO cells could be achieved by compensatory upregulation of other HSPs. Therefore, we looked at levels of mRNA encoding different HSP subgroups that might me upregulated to compensate the loss off HSPA1A/HSPA1B. We found a decrease in HspA8 and HspB1 mRNA levels upon acute MG-stress in HSPA1A/HSPA1B KO cells (Fig S2A,B). In WT cells, mRNA encoding HspB1 and HspA5 was downregulated after short-term MG exposure (Fig S2B). Hsp90AA1 was slightly increased after MG-stress in both, WT and HSPA1A/HSPA1B KO cells. Dnajb1 (DnaJ homolog subfamily B member 1) mRNA was only upregulated HSPA1A/HSPA1B KO cells and HspH1 only in WT cells after acute MG exposure (Fig S2A,B). mRNAs encoding the mitochondrial chaperones HspA9 and HspD1 were both induced upon acute MG-stress in WT as well as in HSPA1A/HSPA1B KO cells (Fig S2C). The co-chaperone of HspD1, Hsp10, showed no changes in mRNA expression upon MG-stress (Fig S2C). mRNA encoding Hsf1 (heat shock factor 1) dropped in HSPA1A/HSPA1B KO cells after short-term MG exposure (Fig S2D).

As mRNA levels encoding HspA9 and HspD1 were induced upon acute MG-stress, we also analyzed HspA9 and HspD1 protein expression after short-term MG exposure. In WT and HSPA1A/HSPA1B KO cells, HspA9 and HspD1 were exclusively expressed in the mitochondria and upregulated upon MG-stress (Fig 2A). However, induction of HspA9 (Fig 2B) and HspD1 (Fig 2C) was higher in HSPA1A/HSPA1B KO cells.

**Fig 2.**
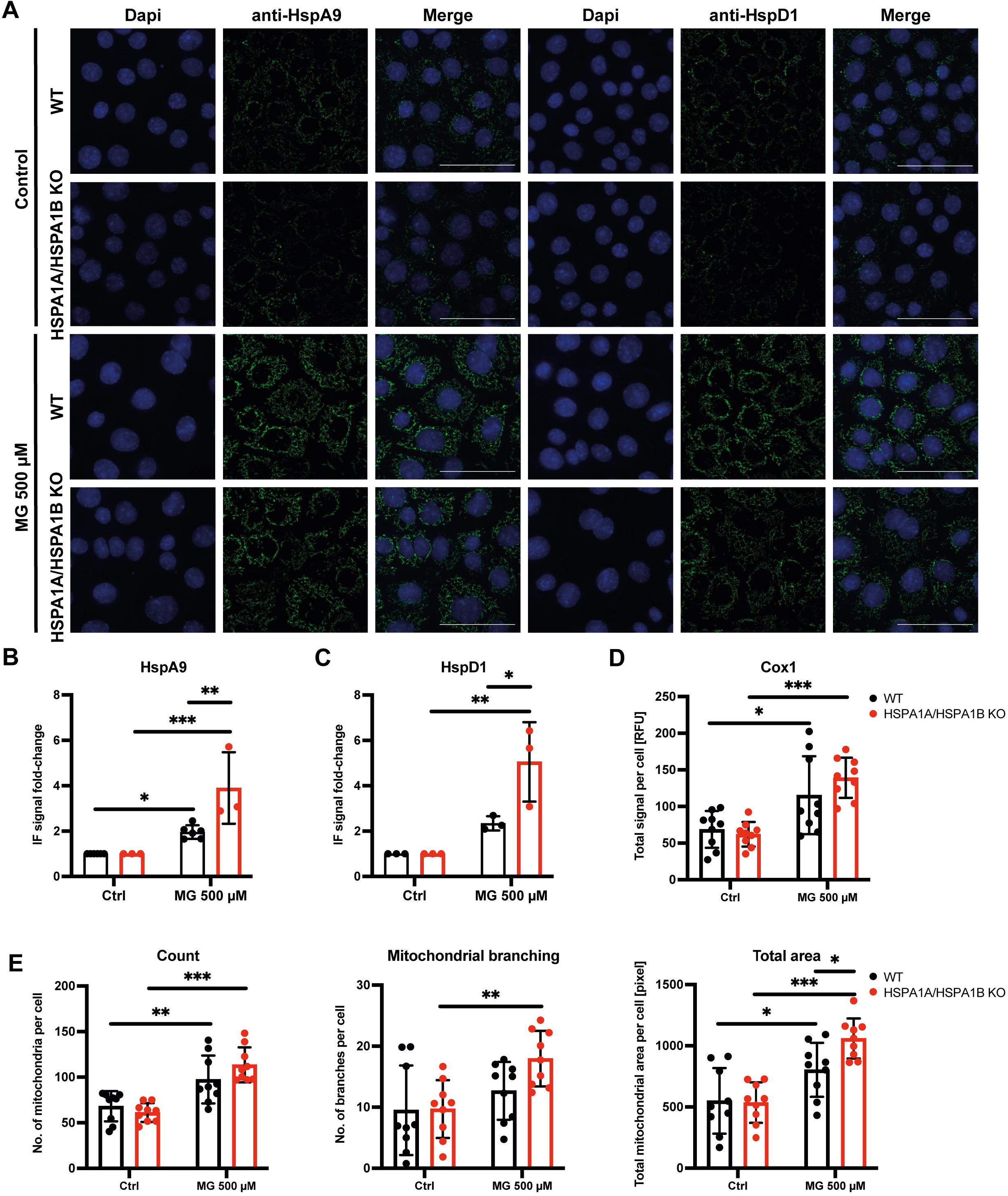
MG-stress induces mitochondrial heat shock proteins and mitochondrial biogenesis. WT and HSPA1A/A1B KO cells were treated with MG for 24 hrs, probed for the mitochondrial HSPs HspA9 and HspD1, analyzed by IF imaging (A) and quantified (B, C). For analysis of changes in mitochondrial morphology upon MG-stress, WT and HSPA1A/A1B KO cells were probed with an anti-Cox1 antibody, imaged via IF, followed by a fully computed analysis of mitochondrial network parameters. (D) Quantification of Cox1 signal. (E) Quantification of mitochondrial count, branching and total area. Results are shown as means ±SD of at least 3 independent experiments. Two-way ANOVA with Šídák’s multiple comparisons test was performed for statistical analysis, *p<0.05, **p<0.01, ***p<0.001.

### 3.3 MG-stress induces mitochondrial biogenesis which is counteracted by HspA1a/HspA1B

The immunofluorescence (IF) stainings of MG-stressed cells suggested changes of mitochondrial network upon MG-stress. Therefore, we complemented IF imaging for Cox1 (cytochrome c oxidase subunit 1) followed by a fully computed mitochondrial network analysis. Total Cox1 signal increased after acute MG-stress and was even stronger in HSPA1A/HSPA1B KO as compared to WT cells (Fig 2D). The same applied for mitochondrial count, mitochondrial branching, and mitochondrial total area (Fig 2E).

### 3.4 Acute MG exposure reduces mitophagy and induces the mitochondrial unfolded protein response

We next questioned the underlying mechanisms leading to induction of HspA9, HspD1 and mitochondrial mass upon MG-stress. Therefore, we looked at key-regulators of mitochondrial fusion, fission, mitophagy and the mitochondrial unfolded protein response (mtUPR). mRNA encoding fusion proteins Mfn2 (Mitofusin-2) and Opa1 (Dynamin-like 120 kDa protein, mitochondrial) were unchanged, whereas mRNA encoding Mfn1 (Mitofusin-1) was downregulated in WT and upregulated in HSPA1A/HSPA1B KO cells upon acute MG-stress (Fig 3A). The fission protein Drp1 (Dynamin-1-like protein) was induced in both cell lines, whereas Fis1 (mitochondrial fission 1 protein) was reduced in both cell lines on mRNA level after short-term MG exposure (Fig 3B). Proteins involved in mitophagy, namely Parkin, Pink1 (PTEN-induced kinase 1) and Bnip3 (BCL2/adenovirus E1B 19 kDa protein-interacting protein 3), were decreased on mRNA level upon MG-stress (Fig 3C). Components of the TIM23 complex, Timm17a (Mitochondrial import inner membrane translocase subunit Tim17-A) and Timm23 (Mitochondrial import inner membrane translocase subunit Tim23), were increased in HSPA1A/HSPA1B KO cells as markers of mitochondrial biogenesis or mtUPR (Fig 3D). Mitokines are known to be changed in mitochondrial stress conditions, therefore we also looked at mRNA levels encoding Fgf21 (Fibroblast growth factor 21) and Gdf15 (Growth/differentiation factor 15). We found that Fgf21 mRNA levels were decreased in both cell lines upon MG-stress, whereas Gdf15 was strongly induced in HSPA1A/HSPA1B KO cells (Fig 3E).

**Fig 3.**
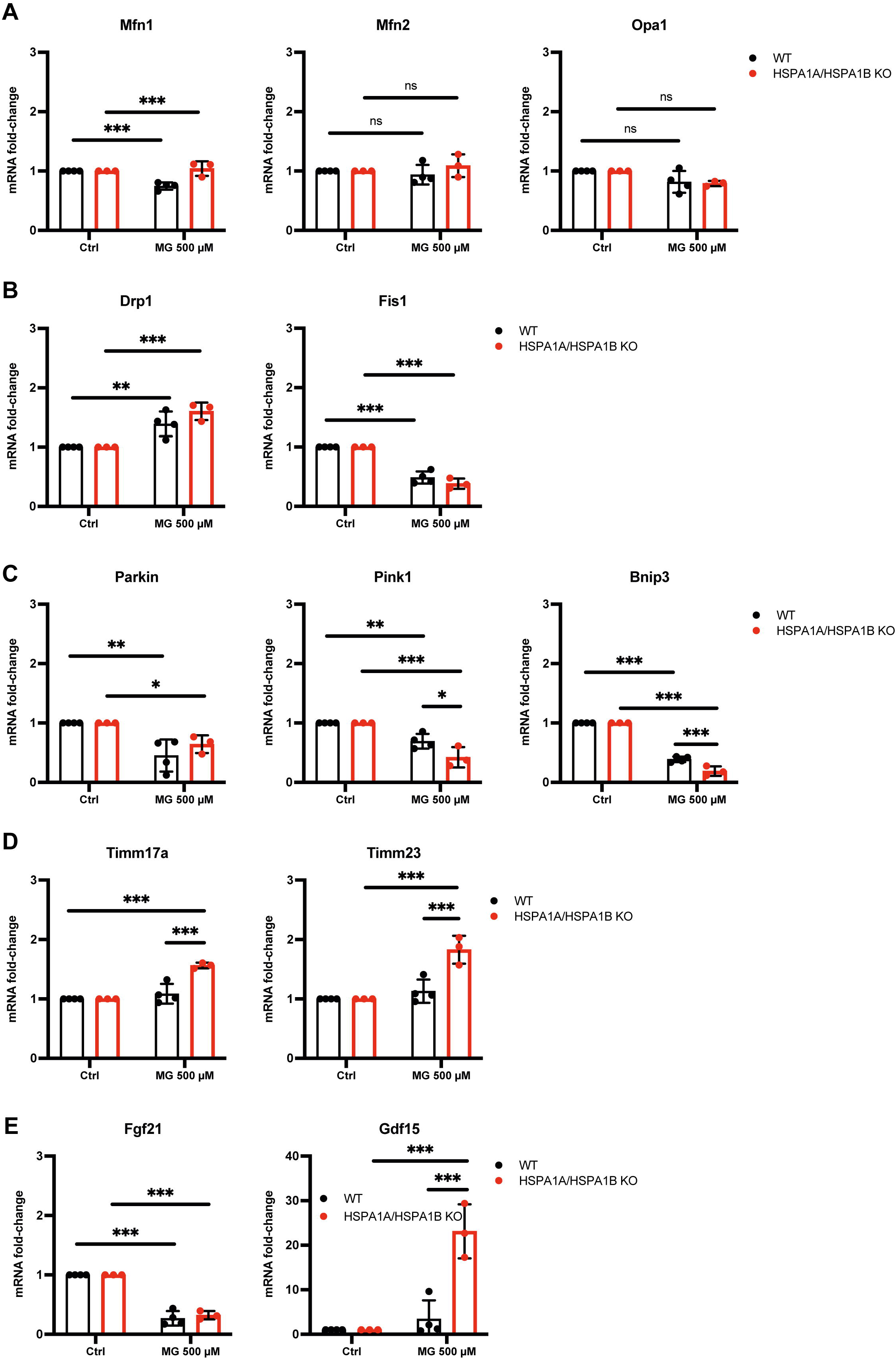
Acute MG exposure reduces mitophagy and induces the mitochondrial unfolded protein response. WT and HSPA1A/A1B KO cells were treated for 12 hrs with 500 µM MG before mRNA analysis. (A) Levels of mRNA encoding fusion proteins Mfn1, Mfn2, Opa1. (B) Levels of mRNA encoding fission proteins Drp1 and Fis1. (C) mRNA expression levels of mitophagy mediators Parkin, Pink1 and Bnip3. (D) mRNA expression of the key components of the TIM23 complex, Timm17a and Timm23. (E) Levels of mRNA encoding the mitokines Fgf21 and Gdf15. Results are shown as means ±SD of at least 3 independent experiments. Two-way ANOVA with Šídák’s multiple comparisons test was performed for statistical analysis, *p<0.05, **p<0.01, ***p<0.001.

### 3.5 Transcription factors involved in mitochondrial biogenesis are predominantly changed in HSPA1A/A1B KO cells upon MG-stress

To identify the transcription factors involved in induction of mitochondrial biogenesis, we looked at mRNA levels of Ppargc1a and Ppargc1b (Peroxisome proliferator-activated receptor gamma coactivator 1-beta), as they are known to be co-regulators of Nrf1 (Nuclear factor erythroid 2-related factor 1) and Nrf2 (Nuclear factor erythroid 2-related factor 2), which promote the expression of Tfam (mitochondrial transcription factor A) (26, 46, 47). We found that Ppargc1a and Ppargc1b are downregulated upon MG-stress in WT cells (Fig 4A). In HSPA1A/HSPA1B KO cells only Ppargc1a mRNA was downregulated (Fig 4A). Nrf2 mRNA levels were unchanged, whereas Nrf1 expression increased in HSPA1A/HSPA1B KO cells but not in WT cells after acute MG exposure (Fig 4B). For transcription of mtDNA, the transcription factors Tfam and Tfb2m (mitochondrial transcription factor B2) are needed (48). Both were significantly upregulated in HSPA1A/HSPA1B KO but not in WT cells upon acute MG-stress (Fig 4C).

**Fig 4.**
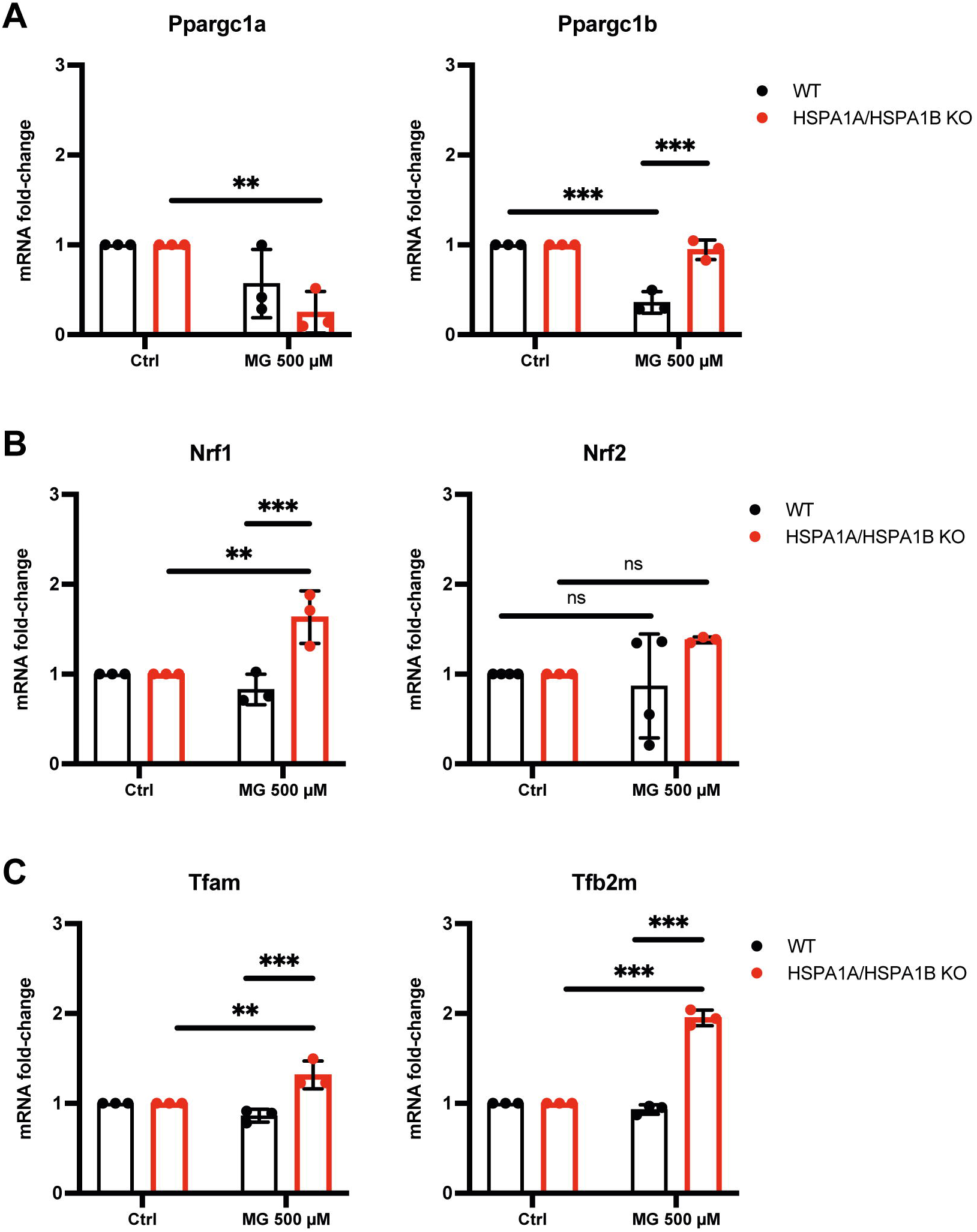
Transcription factors involved in mitochondrial biogenesis are predominantly changed in HSPA1A/A1B KO cells upon MG-stress. Levels of mRNA encoding transcription factors regulating mitochondrial biogenesis were analyzed in WT and HSPA1A/A1B KO cells after 12 hrs incubation with 500 µM MG. (A) mRNA expression of Ppargc1a and Ppargc1a, (B) of Nrf1 and Nrf2, (C) of Tfam and Tfb2m. Results are shown as means ±SD of at least 3 independent experiments. Two-way ANOVA with Šídák’s multiple comparisons test was performed for statistical analysis, *p<0.05, **p<0.01, ***p<0.001.

## 4. Discussion

We found that acute MG-Stress induces the expression of mitochondrial HspA9 and HspD1 as well as mitochondrial biogenesis, both of which are counterregulated by stress-inducible HspA1A/HspA1B. These data suggest a prominent role of tight regulation of mitochondrial biogenesis by HSPs during acute MG-stress.

Increased MG-accumulation in endothelial cells has been linked to induction of the unfolded protein response (49). To show that mitochondrial homeostasis is of importance for securing ATP-dependent processes of the PQC, we knocked out HSPA1A/HSPA1B in mouse cardiac endothelial cells to increase proteotoxic and mitochondrial stress under acute MG exposure. Our hypothesis was, that loss of HSPA1A/HSPA1B would lead to increased MG-H1 accumulation, toxicity, and mitochondrial dysfunction upon MG-stress.

We found that HSPA1A/HSPA1B KO cells accumulate higher MG-H1 levels as compared to WT cells, however MG-H1 clearance was highly efficient in the KO cells after the medium was changed. Indeed, MG-H1 concentrations in HSPA1A/HSPA1B KO cells dropped to the same level as in WT cells after further 24 hrs (Fig 1C). This might explain why we did not observe a significant increase in MG-toxicity in the HSPA1A/HSPA1B KO cells as compared to WT cells (data not shown). To identify compensatory mechanisms that would enable the HSPA1A/HSPA1B KO cells to clear MG-H1, we analyzed the expression levels of other HSPs.

Here we found the mitochondrial heat shock proteins HspA9 and HspD1 to be the strongest induced HSPs upon acute MG-stress, both on mRNA and protein levels (Fig 2A, B, C, Fig S2C). Upregulation of HspA9 and HspD1 was present in WT and HSPA1A/HSPA1B KO cells, however induction on protein level was pronounced in HSPA1A/HSPA1B KO cells upon MG-stress (Fig 2B, C). We first interpreted the induction of HspA9 and HspD1 expression as activation of the mtUPR due to rising proteotoxic stress. Therefore, we hypothesized that acute MG-stress would lead to disruption of mitochondrial homeostasis, which would be pronounced in the absence of HSPA1A/HSPA1B.

However, we found an increase in total mitochondria count, mitochondrial area, and mitochondrial branching (Fig 2E) after acute MG exposure. These changes were again even stronger in HSPA1A/HSPA1B KO cells and accompanied by downregulation of mRNA encoding proteins involved in fission and mitophagy (Fig 3B, C). Overall, these changes suggested that cells initiate mitochondrial biogenesis upon MG-stress, which was generally pronounced in the absence of HSPA1A/HSPA1B. The induction of mitochondrial biogenesis was further supported by the upregulation of transcription factors Nrf1, Tfam and Tfb2m in HSPA1A/HSPA1B KO cells, which are known mediators of mitochondrial biogenesis. We explained the increasing mitochondrial mass as a compensatory cellular mechanism to supply sufficient ATP levels for effective clearance of modified proteins under acute MG-stress. Hence, HSPA1A/HSPA1B KO cells would need stronger induction of mitochondrial biogenesis as they initially accumulate higher MG-H1 concentrations and have a higher demand in preventing aggregation of misfolded and damaged proteins. Therefore, the hormetic autofeedback induced by MG results in acute stimulation of mitochondrial biogenesis which is counterreagulated by HspA1A/HspA1B thereby preventing from mitochondrial overstimulation and exhaustion.

If upregulation of mitochondrial mass would be a central defense mechanism to handle rising proteotoxic stress, induction of HspA9 could also be explained by increased translocation of proteins into the mitochondrial matrix during mitochondrial biogenesis, as it is the central subunit of the PAM (presequence translocase-associated motor) (50). Also, HspA9 could mirror the need for increased oxidative stress defense, as it has been described to play a key role in tumor survival especially under oxidative stress (51). HspA9 has been shown to mediate hypoxia induced preconditioning by preserving the activity of Cox1 and hence decreasing mitochondrial reactive oxygen species (ROS) production (52). Furthermore, downregulation of HspA9 resulted in increased autophagy and decreased pexophagy (53), which might explain why we observed a decrease in mitophagy upon upregulation of HspA9 via MG-stress.

HspD1 together with Hsp10 assists folding of proteins in the mitochondrial matrix including proteins involved in the synthesis of mitochondrial proteins, the respiratory chain and the mitochondrial PQC (54). Induction of HspD1 upon MG-stress could therefore indicate increased mitochondrial biogenesis as well as activation of the mtUPR. Interestingly, heterozygous HspD1 KO mice have been described to exhibit an altered adipose tissue metabolism with mitochondrial dysfunction and altered autophagy as well as local insulin resistance (55). HspD1 could therefore play a central role in maintaining mitochondrial function under increased metabolic stress conditions.

As mitochondrial dysfunction as well as mtUPR have been linked to Fgf21 and Gdf15, we looked at changes on mRNA levels after acute MG exposure. Both have been shown to be elevated in patients with metabolic syndrome and to be attenuated by exercise intervention (56). However, data on Fgf21 and Gdf15 action are still inconclusive as they seem to prolong lifespan in model organisms (57, 58). Here, we found that Fgf21 was downregulated in WT and HSPA1A/HSPA1B KO cells upon acute MG-stress, whereas Gdf15 was dramatically induced in HSPA1A/HSPA1B KO cells. Therefore, both mitokines might signal mitochondrial changes under increased metabolic stress to other tissues possibly mediating adaptive mechanisms.

Taken together we found that acute MG-stress, as observed during increased metabolic flux, leads to induction of compensatory mechanisms, which consist of upregulation of the mitochondrial chaperones HspA9 and HspD1 as well as induction of mitochondrial biogenesis. These defense mechanisms were even pronounced in the absence of HSPA1A/HSPA1B, possibly because HSPA1A/HSPA1B KO cells initially accumulate higher MG-H1 levels upon MG-stress. To provide effective PQC and functioning of the ATP-dependent HspAs, cells must secure oxidative capacity which is maintained through mitochondrial biogenesis. However, mitochondrial dynamics need tight regulation as overstimulation of mitochondrial biogenesis would also increase oxidative stress and disturb mitochondrial homeostasis. Therefore, permanent induction of these compensatory mechanisms might lead to exhaustion and dysbalance of mitochondrial homeostasis as it is observed in diabetic complications. We found that stress-inducible HspA1A/HspA1B counteract this overstimulation of the MG-induced hormetic autofeedback during increased metabolic-stress.

## 5. Conclusion

Acute MG-stress induces mitochondrial HSPs as well as mitochondrial biogenesis in endothelial cells supporting the hypothesis of MG-induced hormesis. Tight regulation by HspA1A/HspA1B counteracts overstimulation of mitochondrial HspA9 and HspD1 as well as mitochondrial biogenesis. Therefore, HspA1A/HspA1B play a central role in maintaining the positive effects achieved by the MG-induced hormetic autofeedback and in prevention of mitochondrial exhaustion. Understanding of the HSP-regulated hormesis effects on mitochondrial content might reveal novel targets for the prevention and treatment of diabetic complications.

## Supporting information

Mitochondrial network analysis

## Author contributions

Conceptualization: J.Z., S.H.; Methodology: R.B., T.F., R.M., M.F.; Investigation: R.B., T.F., C.R.; Writing original Draft: J.Z., R.B.; Writing Review & Editing: T.F., M.M., S.H., J.S.; Funding Acquisition: J.Z.; Supervision: J.Z.

## Acknowledgments

We thank Prof. Holger Lorenz, head of the imaging facility of the ZMBH, who programmed the mitochondrial network plugins.

Furthermore, we want to thank Prof. Peter Nawroth for critical reading and discussion of the manuscript.

## Supporting information

