## Supplementary Figure legends for "Induction of mitochondrial heat shock proteins and mitochondrial biogenesis in endothelial cells upon acute methylglyoxal stress: Evidence for hormetic autofeedback"

**Fig S1.** Characterization of HspA1A/A1B KO MCEC cells. (A) Fold change of mRNA expression of HspA1A and HspA1B after heat shock in WT and 10 different HspA1A/A1B KO clones. (B) Western Blot analysis of WT and different HspA1A/A1B KO clones pre and post heat shock and densitometric quantification. (C) Frameshift mutations in the HSPA1A and HSPA1B gene of clones AD4 and BE12.

**Fig S2.** Changes in mRNA expression of different heat shock proteins and co-chaperones upon acute MG-stress. WT and HSPA1A/A1B KO cells were treated for 12 hrs with 500 µM MG before mRNA analysis. (A) Levels of mRNA encoding cytosolic HSPs HspA8, Hsp90AA1 and DNajb1. (B) mRNA expression of the co-chaperones HspB1, HspH1 and HspA5. (C) Levels of mRNA encoding mitochondrial HSPs HspA9, HspD1 and Hsp10. (D) mRNA expression of Hsf1. Results are shown as means $\pm$ SD of at least 3 independent experiments. Two-way ANOVA with Šídák's multiple comparisons test was performed for statistical analysis, $^{*}p<0.05$, $^{**}p<0.01$, $^{***}p<0.001$.
