## Supplementary figures and images for "Induction of mitochondrial heat shock proteins and mitochondrial biogenesis in endothelial cells upon acute methylglyoxal stress: Evidence for hormetic autofeedback"

### Figure S1

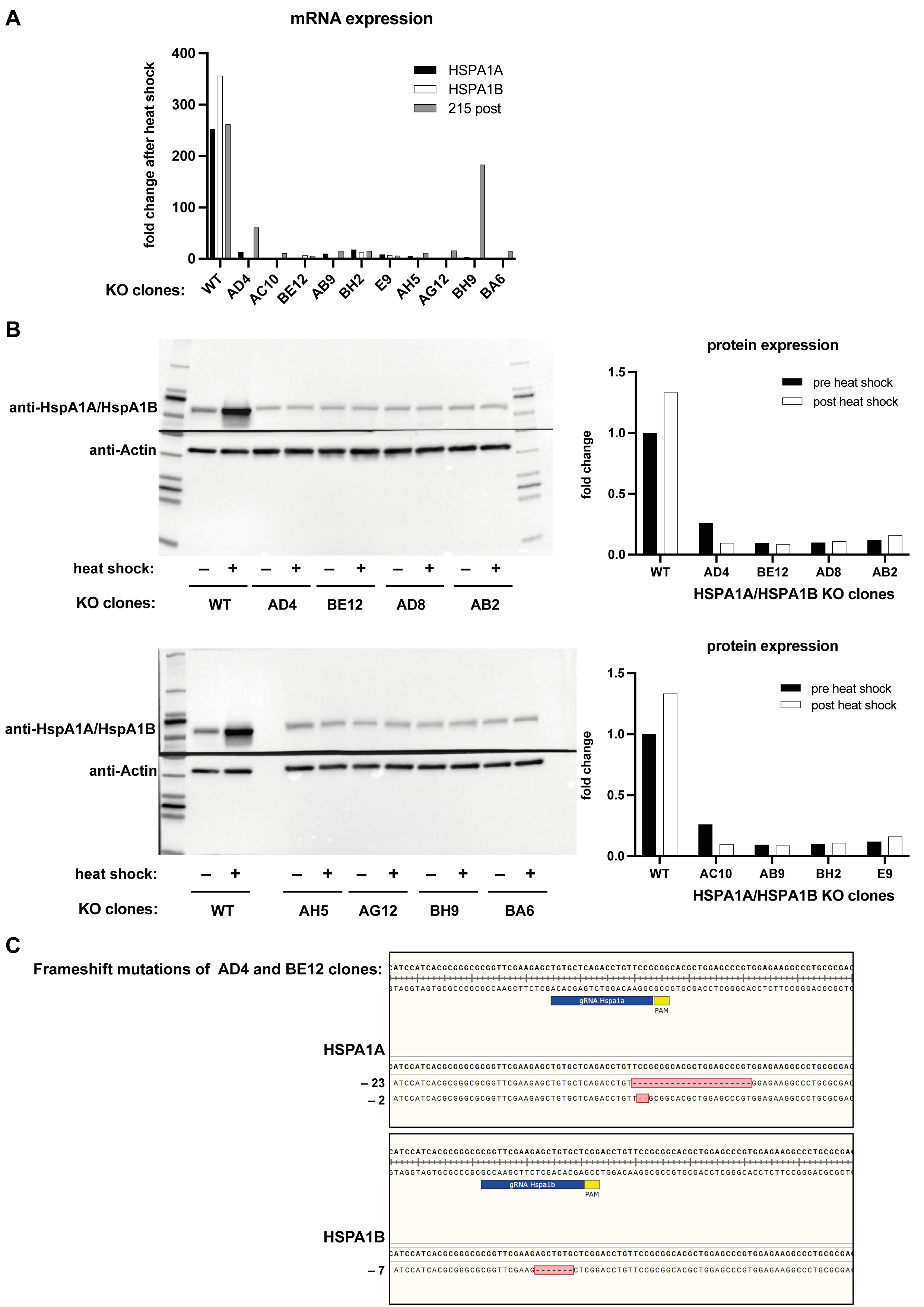

### Figure S2

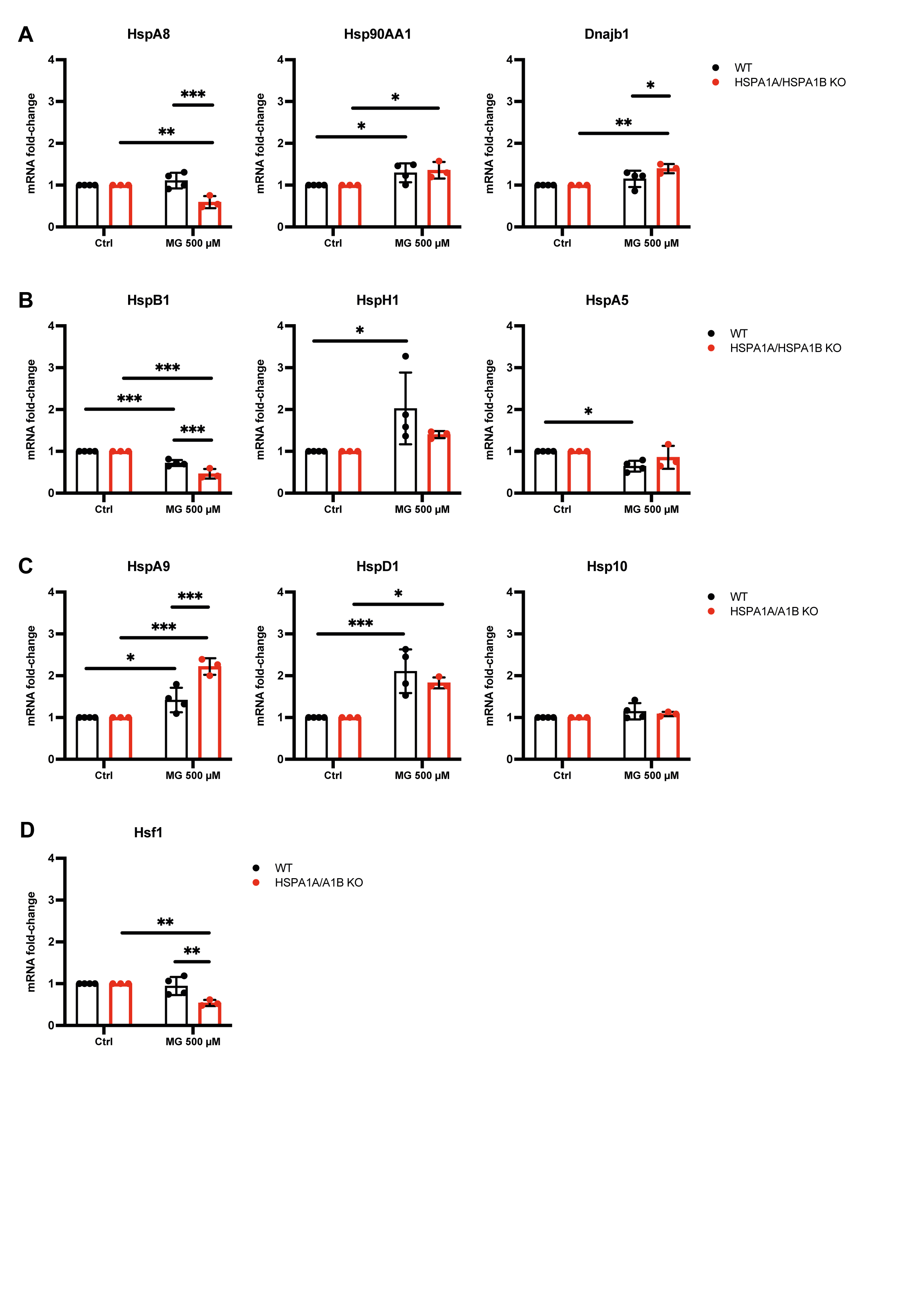
