## Supplementary material for "Induction of mitochondrial heat shock proteins and mitochondrial biogenesis in endothelial cells upon acute methylglyoxal stress: Evidence for hormetic autofeedback": Mitochondrial network analysis

### Mitochondrial network analysis imageJ plugin/macro

Images are acquired as z-stacks of a total of 30 layers with a step size of 200 nm.

Raw image nuclei | Raw image mitochondria

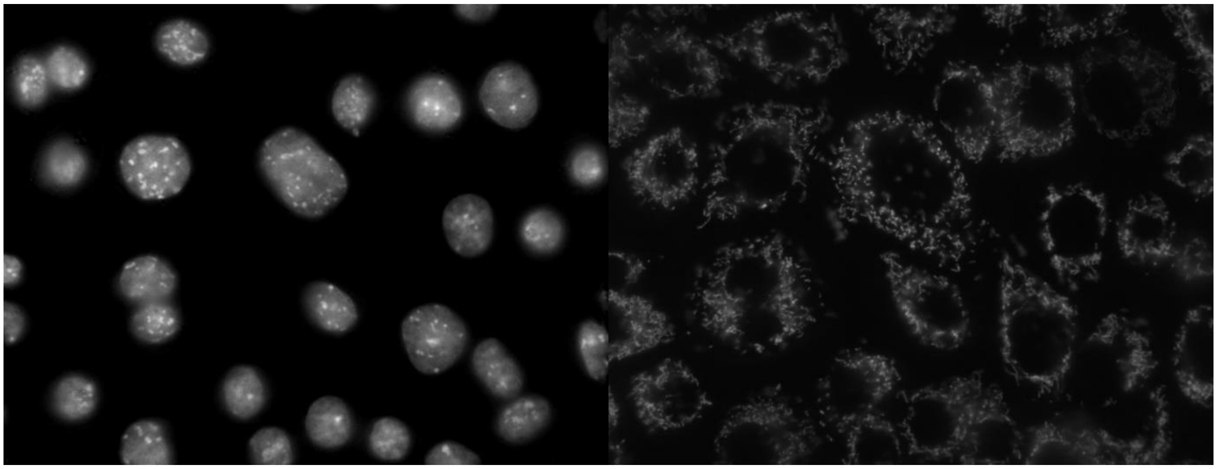

The image stacks of the mitochondrial channels are then deconvoluted using the SVI Huygens Professional software (https://svi.nl/HomePage).

Raw image nuclei | Deconvoluted image mitochondria

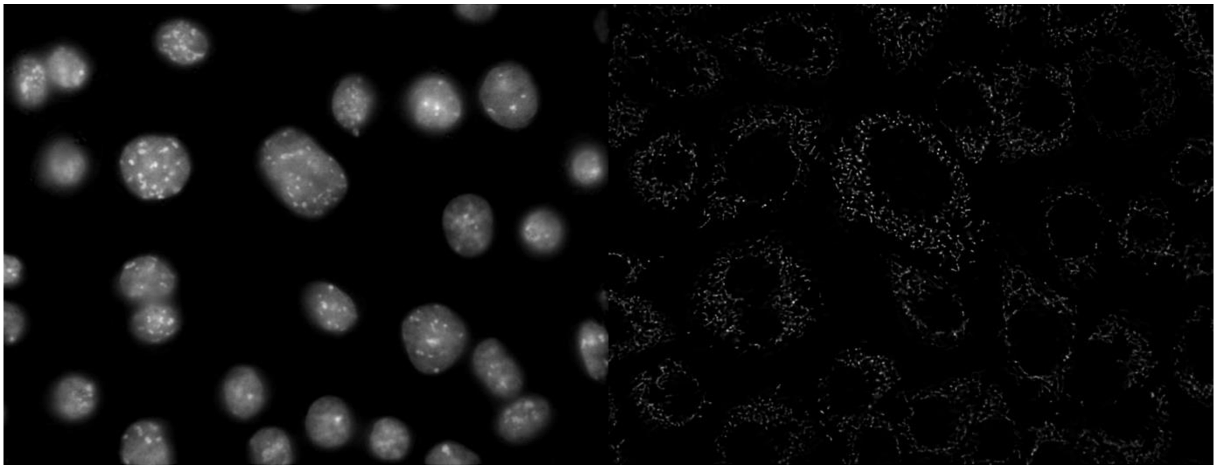

The images are then loaded in imageJ as multicolor image hyperstacks. After running the mitochondrial network analysis plugin/macro, certain options can be set for image processing (background subtraction), thresholding, size and shape discrimination, binary image processing and mitochondria network branching analysis.

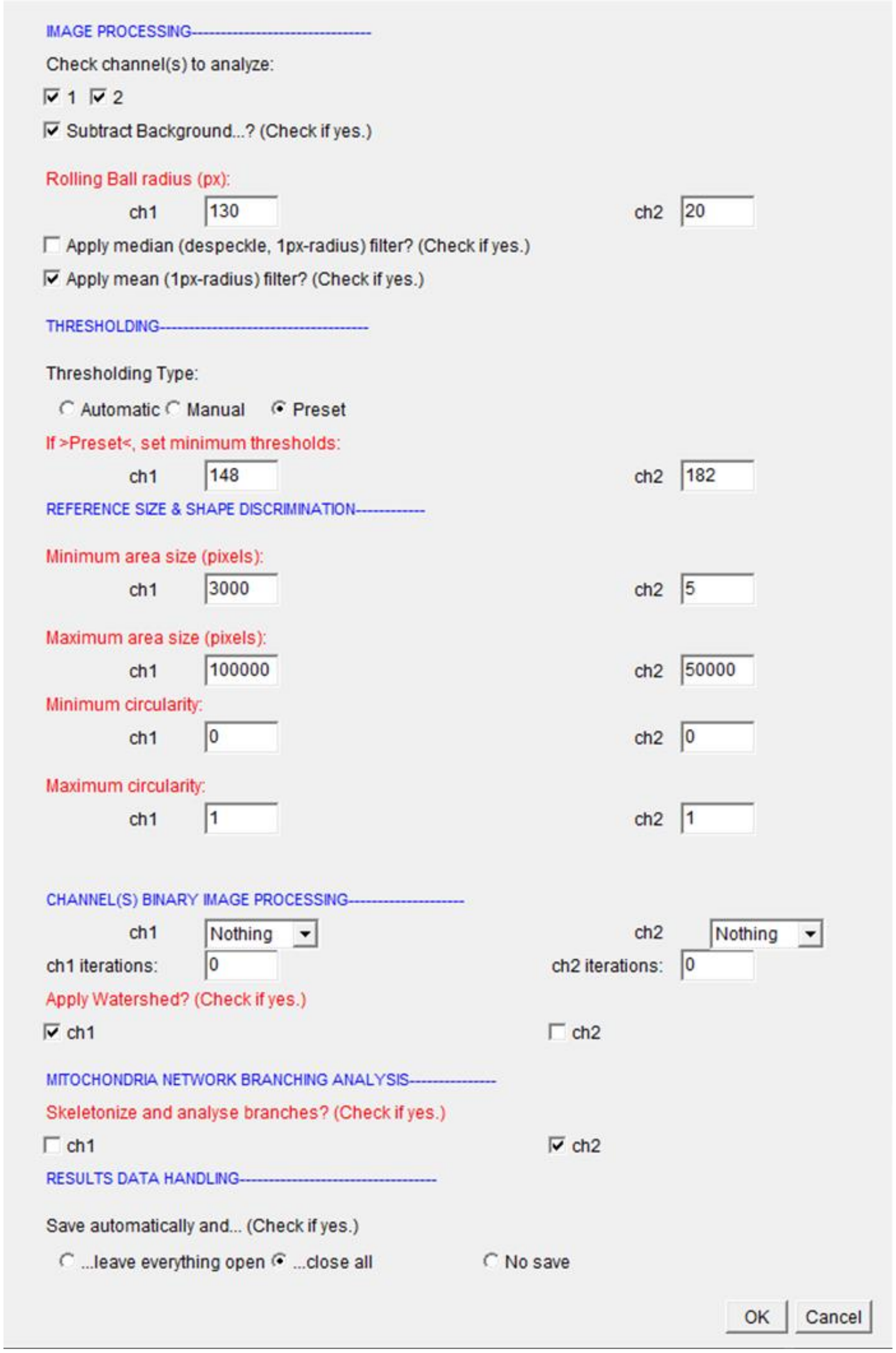


ImageJ processes the images and saves them all separately in a folder. First, the signal is converted into a binary mask. The watershed option can separate two or more nuclei, that are too close to each other and would only appear as one in the binary mask.

Binary mask of nuclei Binary mask of mitochondria

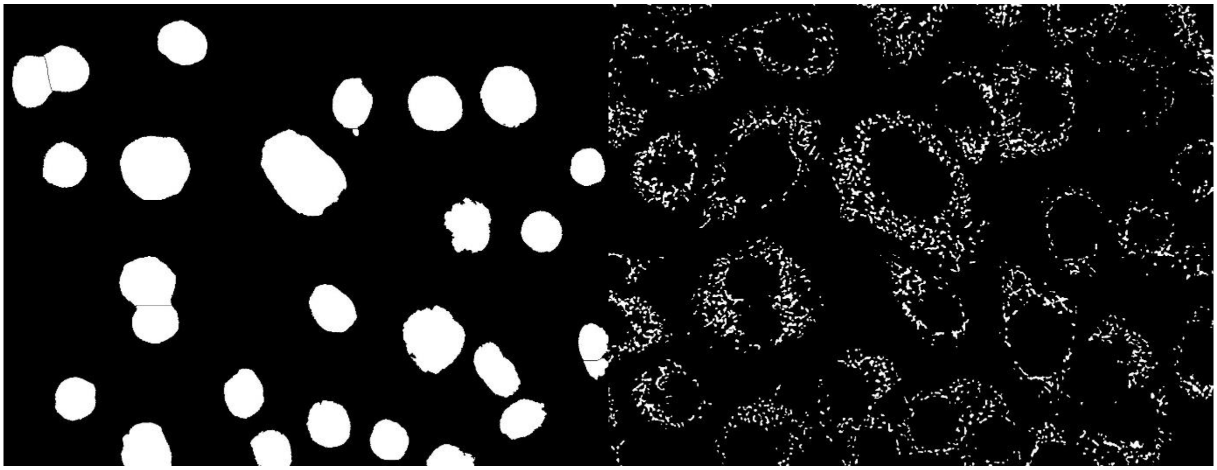

Then the signal from the mitochondria is skeletonized and the information is used for the

- Number of mitochondria
- Total area of mitochondria
- Average size of mitochondria
- % of area
- Mean signal
- Total signal

Skeletonized mask of mitochondria

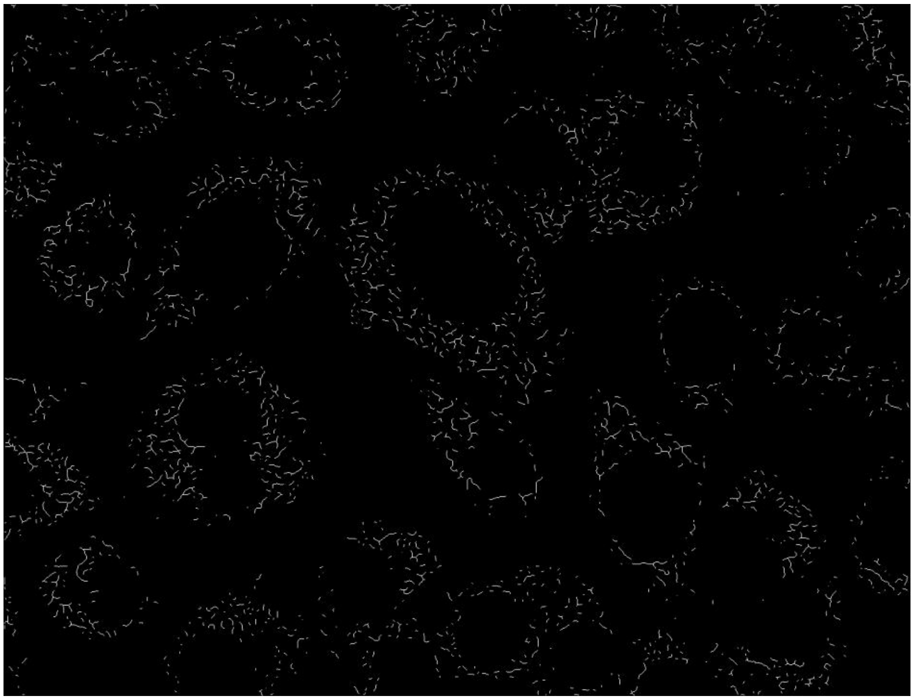

Next, from the skeletonized mask of mitochondria, all locations, which have three or more neighboring pixels are counted as branches/junctions of mitochondria.

Branches/junctions (three or more neighbor pixels)

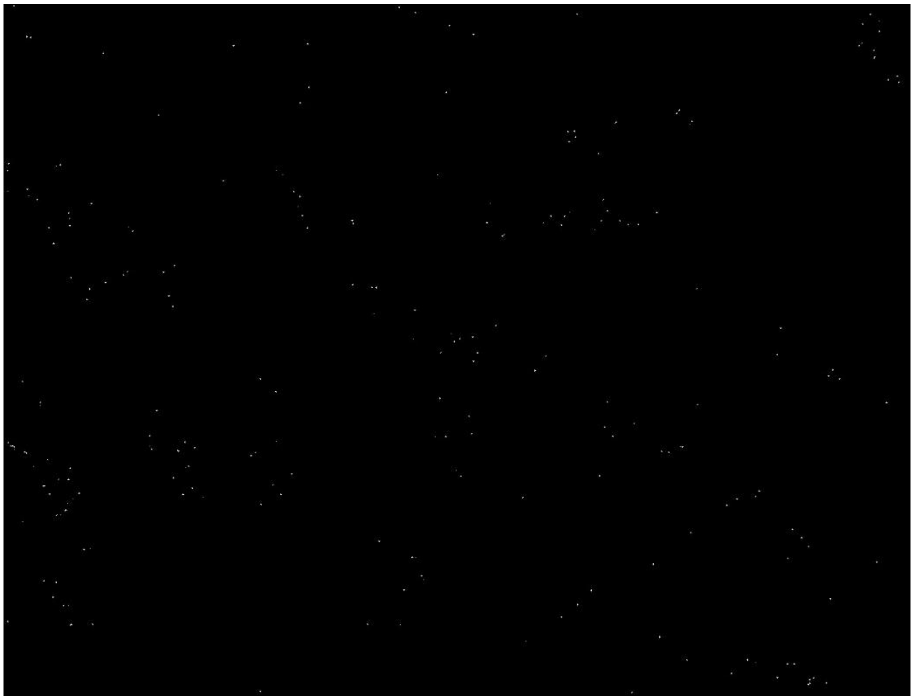

Composite mask of skeletonized mitochondria and junctions

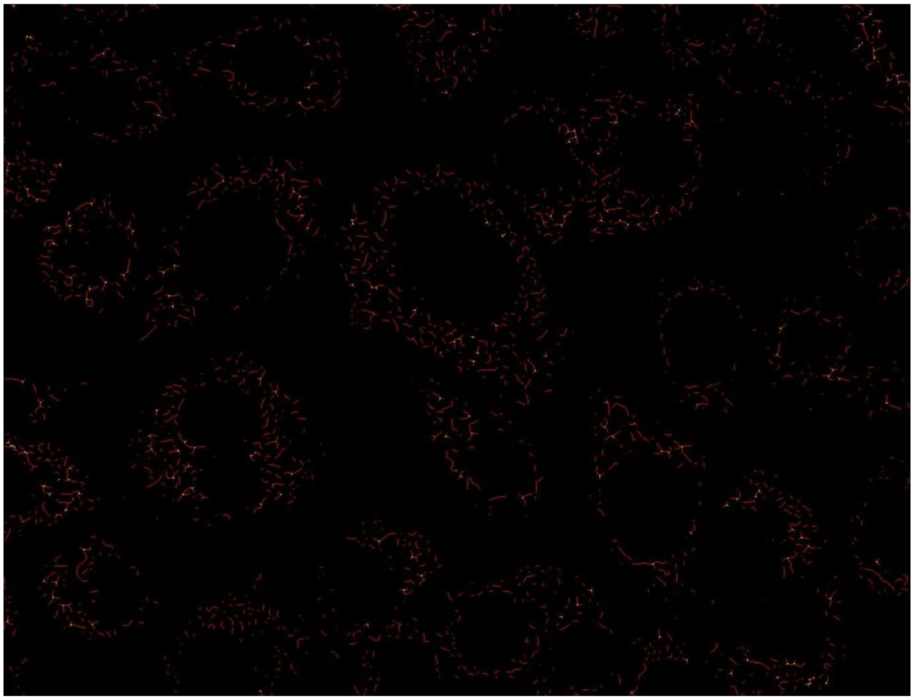
